# Identification and Classification of Expressed Orphan Genes, Spurious Orphan Genes, and Conserved Genes in the Human Gut Microbiome

**DOI:** 10.64898/2026.01.16.699869

**Authors:** Chen Chen, Nikolaos Vakirlis, Rens Holmer, Dick de Ridder, Anne Kupczok

## Abstract

Orphan genes - genes lacking detectable homologs outside a species - are widespread in microbial genomes and are thought to contribute to their adaptation and molecular innovation. However, not all predicted orphan genes may represent novel functional coding sequences. False positive orphan genes, also called spurious orphan genes, can arise from gene prediction errors. We reason that orphan genes lacking detectable expression are more likely to be spurious. To test this, we combined large-scale metatranscriptomic profiling of the human gut microbiome with machine learning to distinguish expressed orphan genes from spurious ones and to compare them with conserved genes found in multiple species. Using nearly 5,000 metatranscriptome libraries, we identified ∼218,000 orphan genes supported by expression evidence, while ∼330,000 predicted orphan genes lacked detectable expression, and were classified as spurious. We extracted 154 features for sequence, structural, and evolutionary properties for each gene and trained XGBoost classifiers while accounting for genomic representation. The models achieved an area under the receiver operating characteristic curve (AUC) of 0.82 in distinguishing expressed orphan genes from spurious orphan genes and an AUC of 0.93 in distinguishing expressed orphan genes from conserved genes. SHAP-based interpretation revealed clear biological signals. Particularly, expressed orphans were present in more genomes than spurious ones and expressed orphan genes were shorter than conserved genes. This work improves orphan gene discovery and suggests that expressed orphan genes differ systematically from conserved genes and spurious orphan genes in sequence composition, structural constraints, and evolutionary signals.

**Significance statement:** Species-specific genes, also called orphan genes, are abundant in bacterial genomes and have the potential to drive evolutionary innovation. However, the study of orphan genes is confounded by annotation errors that produce spurious gene predictions. Here, we leveraged nearly 5,000 metatranscriptomes from the human gut microbiome and implemented machine learning models to distinguish expressed orphan genes from non-expressed, likely spurious, genes and conserved genes. We show that a substantial fraction of predicted orphan genes lacks expression support and they exhibit distinct sequence and evolutionary signatures from expressed orphans. By integrating expression evidence with sequence-, structure-, and evolution-based features, our work refines orphan gene catalogs. This work improves the reliability of orphan gene discovery and provides new insights into the biological characteristics of orphan genes in human microbial communities.

## 1. Introduction

Orphan genes (OGs), also known as orphans, are defined as having no homologs outside a given species or outside a particular lineage (Dujon 1996; Domazet-Loso and Tautz 2003; Li et al. 2022). OGs are ubiquitous in each newly sequenced genome, including those of bacteria, archaea, and phages (Tautz and Domazet-Lošo 2011). Compared to conserved genes (CGs), which occur in multiple species, OGs are characterized by fewer exons in eukaryotes, shorter lengths, higher Adenine-Thymine (AT) content, higher disorder protein structure, higher nonsynonymous substitution rate, and higher isoelectric points (Daubin and Ochman 2004; Fakhar et al. 2023; Vakirlis and Kupczok 2024).

OGs are thought to play a significant role in evolution and speciation, likely contributing to crucial species- or lineage-specific adaptive processes by facilitating the generation of novel functions (Tautz and Domazet-Lošo 2011; Arendsee et al. 2014; Vakirlis et al. 2020; Fakhar et al. 2023; Vakirlis and Kupczok 2024). However, the role of OGs in evolution remains elusive due to their lack of phylogenetic conservation and their origination by diverse of evolutionary processes (Toll-Riera et al. 2009; Tautz and Domazet-Lošo 2011; Fakhar et al. 2023). OGs potentially arise de novo from non-coding genomic regions or alternative reading frames of existing genes, through fast divergence, or by horizontal gene transfer (HGT) (Tautz and Domazet-Lošo 2011; Vakirlis and Kupczok 2024). Thus, OGs originate by different mechanisms and are of different age, forming a heterogenous group (Pereira et al. 2025).

Some experimental studies have shown that proteins derived from random sequence libraries can acquire biologically relevant functions in prokaryotic systems (Babina et al. 2023; Frumkin et al. 2025). This suggests that even short, newly emerged sequences may rapidly become functional, supporting the idea that orphan genes can contribute to evolutionary innovation. Most of our current understanding of OGs and the evolutionary mechanisms has been derived from studies in eukaryotes (Tautz and Domazet-Lošo 2011; Prabh and Rödelsperger 2019). In addition to studies in eukaryotes, recent work has highlighted the importance of orphan genes in non-eukaryotic systems, including viruses, where lineage-specific genes can play key roles in host interaction and adaptation (Nomburg et al. 2024). Additionally, studying prokaryote orphan genes is particularly interesting and relevant for a number of reasons (uz-Zaman and Ochman 2025). Prokaryotes are ubiquitous, ample genome data is available, and they are found in all ecosystems and play crucial roles in various ecological processes, such as nitrogen fixation and organic matter decomposition, which are essential for ecosystem functioning and stability (Whitman et al. 1998; Meng et al. 2022). Moreover, pathogenic prokaryotes can cause diseases in humans, animals, and plants and there is great interest in determining factors involved in pathogenicity (Zhou et al. 2023). Thus, understanding prokaryotic OGs can provide insights into their unique environmental adaptations and mechanisms of pathogenicity, potentially leading to new treatments and preventive measures.

In a previous study, we constructed an orphan reference dataset of predicted species-specific OGs from the human gut environment — a well-explored habitat accommodating numerous prokaryotic species (Vakirlis and Kupczok 2024). A key question is whether this dataset truly contains only novel, functional coding sequences unique to each species. False positive OGs could arise in two ways. First, non-existent genes may be incorrectly annotated as coding sequences because OGs are typically short and often lack evolutionary conservation (Yu and Stoltzfus 2012); we refer to these as spurious OGs (SOGs). Second, genes that are actually conserved may be misclassified as OGs when distant homologs remain undetected, leading to an overestimation of lineage-specific genes (Weisman et al. 2020). In this study, we focus on the first category, SOGs. Our identification of SOGs is based on the expectation that such spurious predictions lack detectable expression. To collect expression evidence for OGs, we used human gut metatranscriptomics data to distinguish expressed OGs (EOGs) from SOGs, defining EOGs as genes with sufficient coverage by mapped reads across multiple samples. Genes without any expression are labelled SOGs and may be artifacts.

We then classified genes as EOGs and SOGs using machine learning (ML) models. ML offers a powerful tool for classification tasks by effectively learning the relevant features in the data and using them to make accurate predictions. These models, such as random forest (Breiman 2001) and XGBoost (Chen and Guestrin 2016), can be trained to identify complex relationships between input features and output labels. Sequence analysis has been the primary source for studying genes being CGs or OGs, focusing on features such as gene length, GC content, isoelectric point, and codon adaptation index (Vakirlis and Kupczok 2024). Structural properties also differed, particularly protein disorder levels were observed to be higher in OGs (Basile et al. 2017). Recently, accurate protein language models such as ESMFold (Lin et al. 2023) have been proposed, greatly advancing the accuracy of protein structure prediction. Because orphan proteins lack homologous sequences, structure prediction methods relying on multiple sequence alignments (MSAs) often perform poorly. In contrast, protein language model–based approaches have been shown to outperform such methods on orphan proteins (Michaud et al. 2022). Here, we used ProstT5 (Heinzinger et al. 2024), which encodes protein sequence information directly from protein sequences without relying on MSAs, to extract so-called 3Di token representations of protein structures from amino acid sequences. While this approach enables structure-related features to be derived without relying on homologous sequences, it may also introduce biases, as protein language models such as ProstT5 are trained predominantly on datasets enriched for conserved proteins. Consequently, structural representations for orphan proteins, which typically lack homologs, may be less reliable. In this study, we address the following research questions. First, what proportion of predicted orphan genes in the human gut microbiome are likely to be spurious due to annotation artefacts? Second, which features are most informative for distinguishing EOGs from SOGs and which features distinguish EOGs from CGs? To answer these questions, we developed machine learning models using sequence-, evolution-, and structure-based features, and evaluated their performance and feature contributions.

## 2. Results

### 2.1 A large proportion of orphan genes are likely spurious

In earlier work, we found 631,104 unique species-specific prokaryotic orphan genes from the human gut microbiome (Vakirlis and Kupczok 2024). To investigate which of these genes are supported by transcriptional evidence, we retrieved 4,969 prokaryotic human gut metatranscriptomics libraries from the Sequence Read Archive (SRA) (Leinonen et al. 2011) and mapped them to our orphan reference dataset (Figure 1). For each gene, we calculated expression coverage across all libraries, assuming that expressed genes are likely true genes and non-expressed genes may be spurious. A gene with no coverage in all libraries was classified as a spurious orphan gene (SOG). We found that the number of genes detected in at least one library decreased continuously with the coverage cutoff (Figure S1A) and additionally the number of detected genes decreased with the number of libraries in which each gene was detected as expressed (Figure S1B). In the following, we denote a gene with coverage ≥ 1.2 in at least 2 libraries as an expressed orphan gene (EOG). These thresholds were selected to minimize technical artifacts by requiring both a reliable coverage level and consistent detection across multiple libraries.

**Figure 1.**
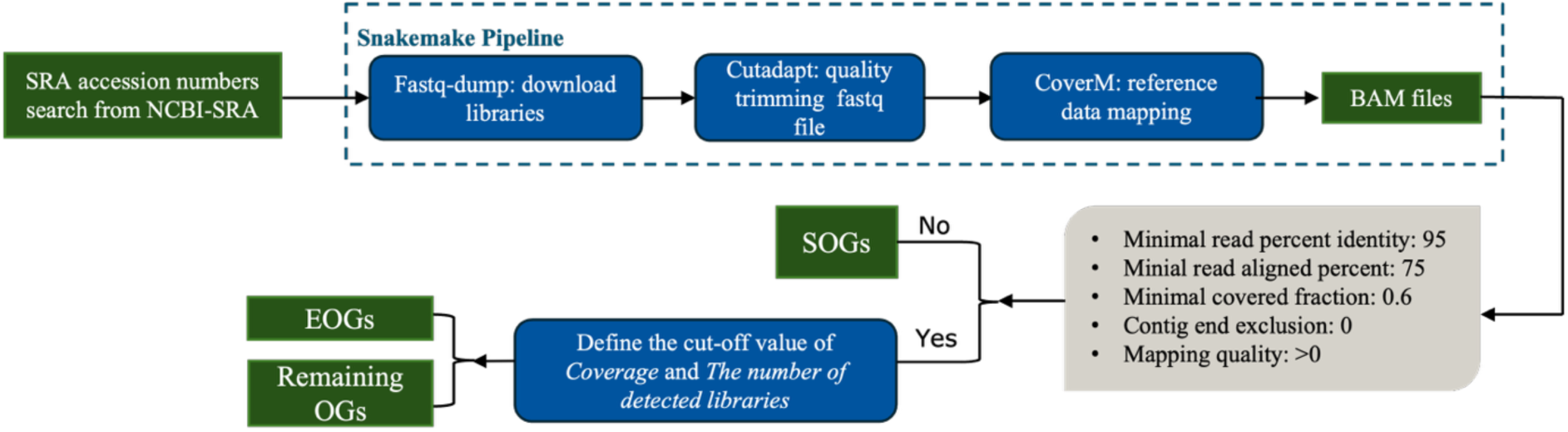
The pipeline used to distinguish spurious and expressed orphan genes.

This classification framework is based on the expectation that genuine protein-coding genes are more likely to exhibit transcriptional evidence than spurious gene predictions. To evaluate whether this expectation is reasonable, we compared expression support between OGs and CGs, where CGs were defined as genes present in more than 10 species. We selected three phylogenetically distinct species with varying numbers of genomes from the subset of species represented by at least 700 genomes in UHGG (Table 1). This threshold was chosen to minimize the effect of the number of genomes in the species on expression detection (see Section 2.2). Crucially, in all three species, CGs were found to be expressed at very high percentages supporting expression as a characteristic of a true gene (Figure 2A). What’s more, CGs consistently showed substantially stronger transcriptional support than OGs: for example, in *Holdemanella* sp002299315, 78.1% of CGs were classified as expressed, whereas only 31.9% of OGs showed detectable expression, and 54.6% of OGs lacked detectable expression entirely. Nevertheless, we observed variation in the number of expressed OGs in this small sample of three species, suggesting that they represent a heterogeneous group with high variation in levels of transcriptional support across species.

**Figure 2.**
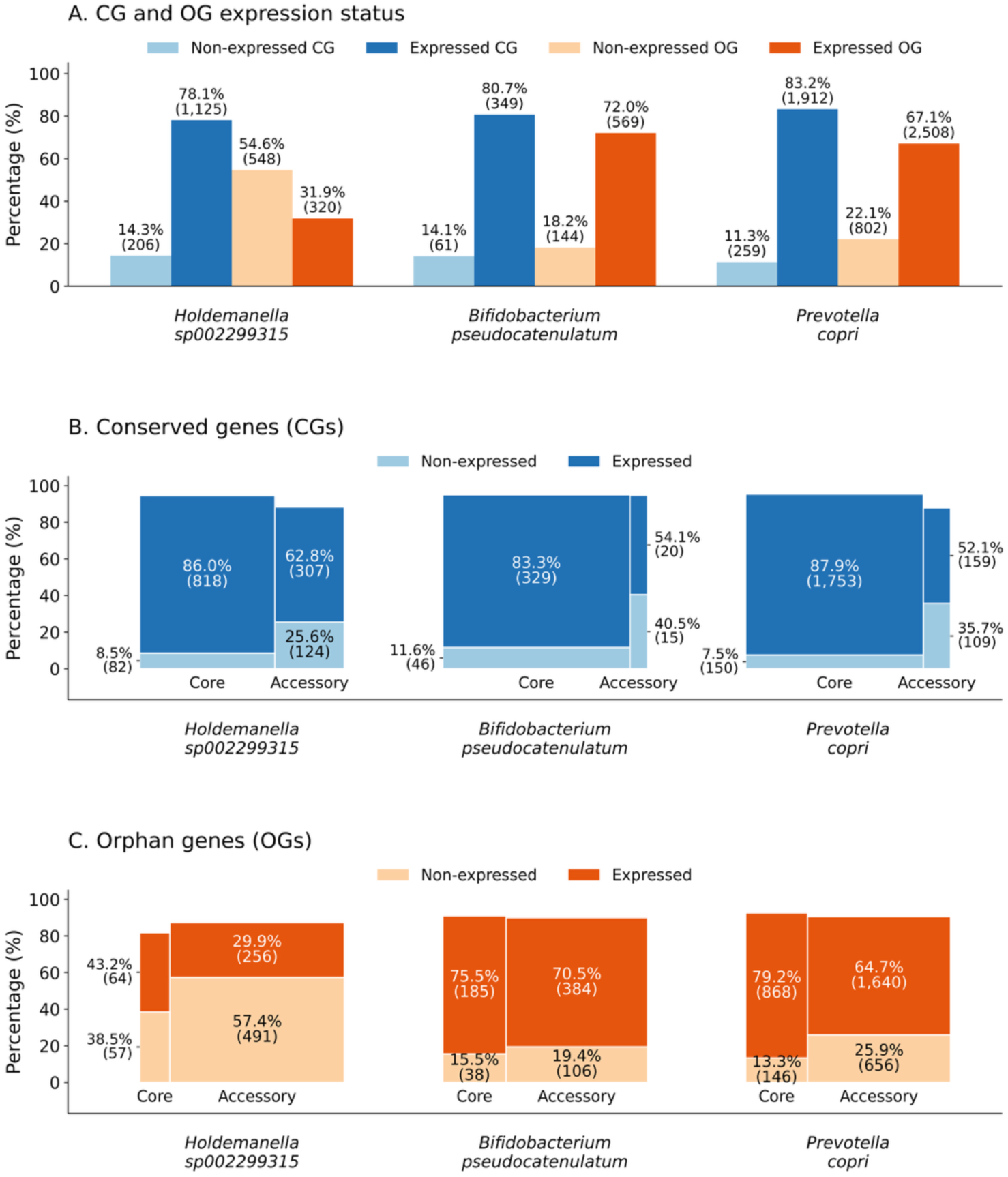
(A) Comparison of expression support between conserved genes and orphan genes across three species. Expression support in core and accessory genes across (B) conserved and (C) orphan genes across three species.

**Table 1.** Representative species selected for expression support analysis.

| Species ID | Number of genomes | Species name |
| --- | --- | --- |
| GUT_GENOME001602 | 995 | *Holdemanella* sp002299315 |
| GUT_GENOME239670 | 1117 | *Bifidobacterium pseudocatenulatum* |
| GUT_GENOME140299 | 2478 | *Prevotella copri* |

To further investigate the heterogeneity of orphan genes, we separated both CGs and OGs into core and accessory categories within each species (Figure 2B,C). Across all three species, the majority of CGs belonged to the core genome (Figure 2B), whereas OGs were predominantly accessory genes (Figure 2C). Core CGs consistently exhibited stronger expression support than accessory CGs. For example, in *Holdemanella* sp002299315, 86.0% of core CGs were expressed compared to 62.8% of accessory CGs, while lack of expression evidence was more frequent among accessory CGs (25.6%) than core CGs (8.5%) and similar trends were observed in the other two species. A similar pattern was observed for OGs. Core OGs generally showed higher expression support than accessory OGs. For example, in *Holdemanella* sp002299315, 43.2% of core OGs were expressed, compared to only 29.9% of accessory OGs. Interestingly, *Holdemanella* sp002299315 also exhibited the lowest proportion of core OGs among the three species and simultaneously showed the lowest overall proportion of expressed OGs. This pattern suggests that species whose OG repertoires are dominated by accessory genes may show lower overall expression support for OGs. Since accessory genes are often low in frequency and strain-specific within species, they may be less consistently detected across metatranscriptomic samples. In addition, previous pangenome studies have shown that low-frequency genes are more enriched in annotation errors or assembly artifacts (Tonkin-Hill et al. 2020), which could contribute to their lower expression support.

Together, these results suggest that the heterogeneous expression patterns of OGs are closely linked to their distribution within the pangenome. Genes belonging to the accessory genome are more likely to lack detectable expression, whereas core genes exhibit substantially stronger transcriptional support. As conserved genes are enriched in core genes, their expression can typically be detected. However, orphan genes contain a higher fraction of accessory genes and the difference in expression evidence between core and accessory genes is less pronounced. This supports our approach to include additional expression evidence for distinguishing EOGs from likely spurious OGs.

Applying the expression-based framework to all OGs resulted in 217,680 EOGs out of the 631,104 reference OGs. A total of 337,985 genes showed no detectable expression and were classified as SOGs. After removing genes with missing feature values and genes originating from archaea, the final dataset contained 214,356 EOGs and 329,765 SOGs. We further examined the robustness of this classification using a rarefaction analysis of metatranscriptome libraries (Figure 3). The number of detected EOGs increased steadily as additional libraries were included, whereas the number of detected SOGs decreased. The EOG curve increased more slowly with larger sample sizes, suggesting that most OGs with readily detectable expression had already been captured and that additional libraries contributed progressively fewer newly detected EOGs. At the same time, the decreasing number of SOGs indicates that some genes initially lacking expression support gain transcriptional evidence as sampling effort increases. Nevertheless, some orphan genes may only be expressed under rare environmental or physiological conditions and may therefore remain undetected in the current dataset. Together, these results suggest that classification depends on the accumulation of expression evidence but approaches stability with increasing sampling effort.

**Figure 3.**
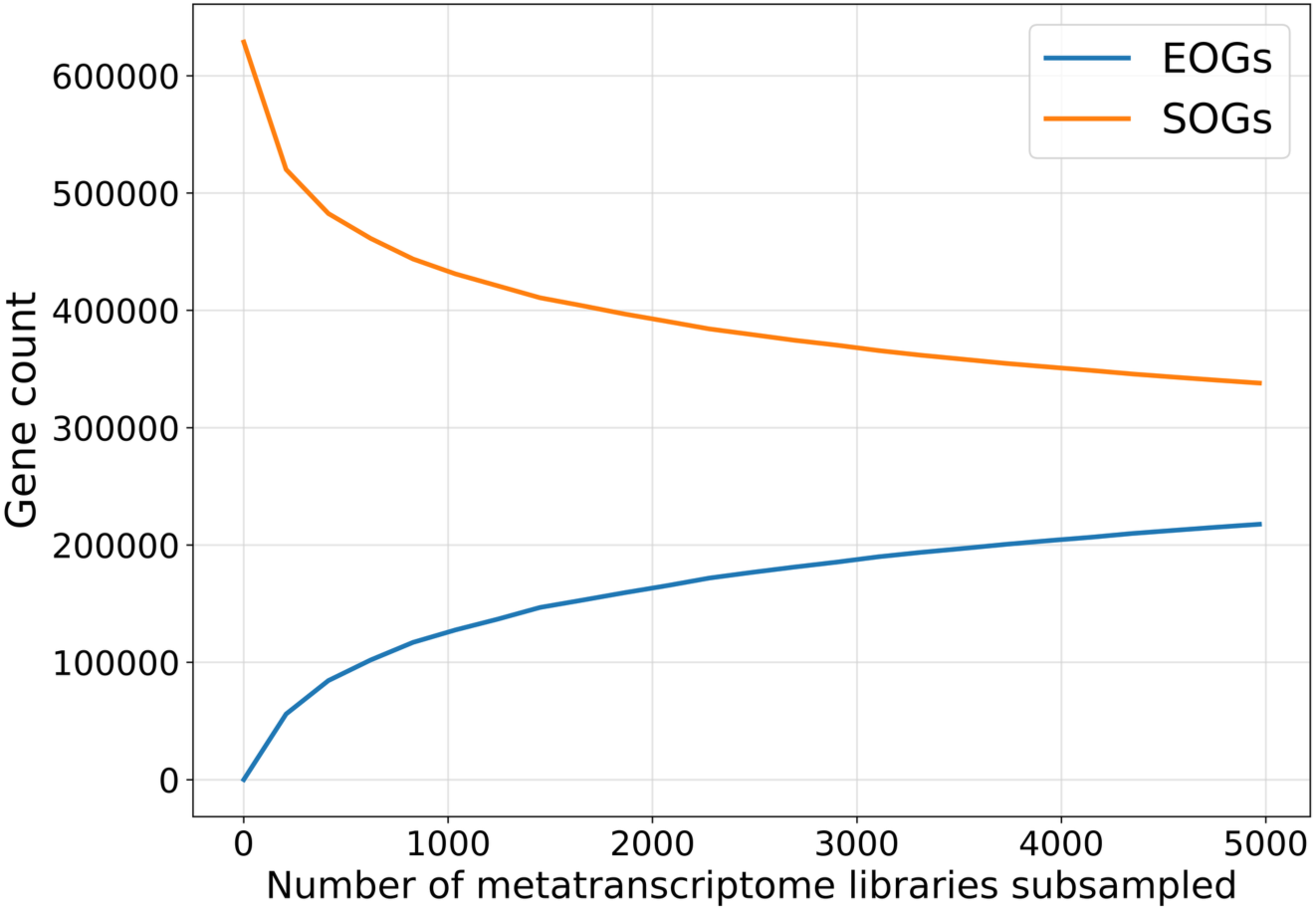
Rarefaction analysis of OG expression. Gene count provides the average number of genes detected among 50 resamplings.

### 2.2 Spurious orphans have distinct properties compared to expressed orphans

We next assessed the ability of models trained on different feature sets to distinguish EOGs from SOGs using a balanced dataset of 214,356 samples per class. We evaluated four groups of features: sequence features (*F_seq_*), structural features (*F_struct_*), evolutionary features (*F_evo_*), and a combined feature set (*F_all_*). Detailed descriptions of these features are provided in the Materials and Methods section. Model performance was evaluated using 5-fold stratified cross-validation, where in each fold approximately 171,485 samples per class were used for training and 42,871 per class for testing, with class balance preserved in all splits. We found that the model based on *F_evo_* features substantially outperformed those based on *F_seq_* and *F_struct_* in distinguishing EOGs from SOGs (Table 2, first row). To understand why this was the case, we examined feature importance and found that the total number of genomes available for the corresponding species (*F_evo_* feature “Number_genomes_in_species”) emerged as the most important feature for classification (Figure S2). This result is supported by the observation that EOGs tend to be found in species represented by a larger number of genomes (Figure S3A). In contrast, CGs and SOGs exhibited similar distributions, with many genes derived from species represented by fewer genomes. The higher genomic representation likely contributes to the predictive power of evolutionary features in distinguishing EOGs from SOGs and might introduce a bias in our analyses.

**Table 2.**
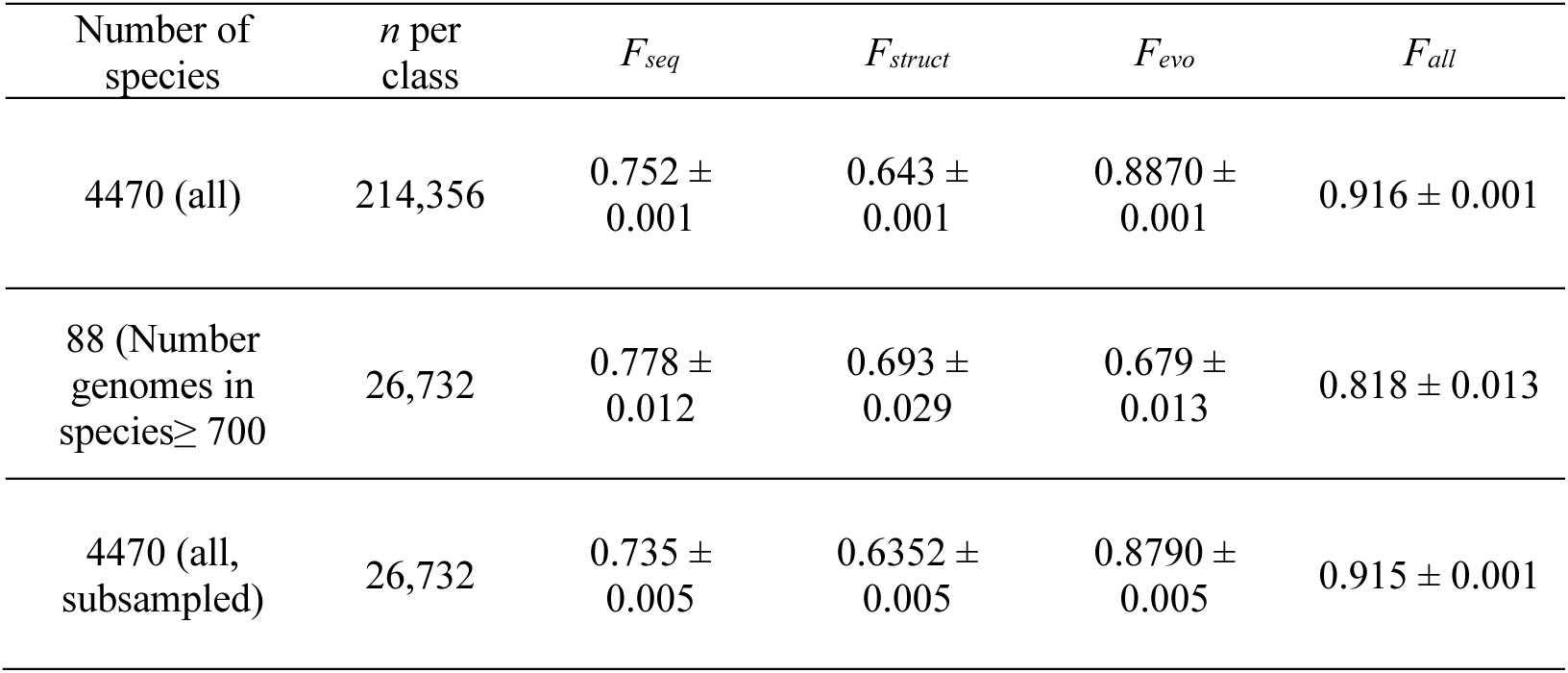
Classification performance (average AUC ± SD across inner folds) for EOGs versus SOGs with various feature sets, shown before (first row, third row) and after (second row) controlling for genomic representation bias.

To address this potential bias introduced by genomic representation, we restricted the analysis to 88 species for which at least 700 genomes were available, resulting in 72,828 EOGs and 26,732 SOGs (Number_genomes_in_species ≥ 700) (Table S1). Under this controlled scenario, the distributions of Number_genomes_in_species between EOGs and SOGs become more similar (Figure S3B), allowing for a less biased evaluation of evolutionary features. Classification after bias correction and using datasets of equal size (26,732 samples per class; 21,386 for training and 5,346 for testing) showed that *F_seq_* yields better performance than *F_evo_* and *F_struct_* (Table 2, second row). Thus, under these conditions, *F_evo_* no longer stands out as providing best performance in the individual feature classes. Additionally, to assess whether the lower mean standard deviation (SD) in the uncorrected analysis (Table 2, first row) was driven by sample size, we repeated the analysis using subsampled datasets matching the size of the bias-corrected set analysis (Table 2, third row). Although the SD increased slightly, it remained substantially lower than that observed after bias correction, indicating that the reduced variance is not primarily attributable to sample size differences, but rather to the presence of the strong representation bias signal, which makes the classification task easier and more stable across folds.

To better understand the protein and gene characteristics that distinguish EOGs from SOGs after controlling for genomic representation bias, we examined the contribution of features to classification performance using SHAP values (Figure 4). We focused on the top 25 features, as individual feature contributions dropped below 1% after the 25th ranked feature, consistent with a long-tail distribution of feature importance (Figure S4).

**Figure 4.**
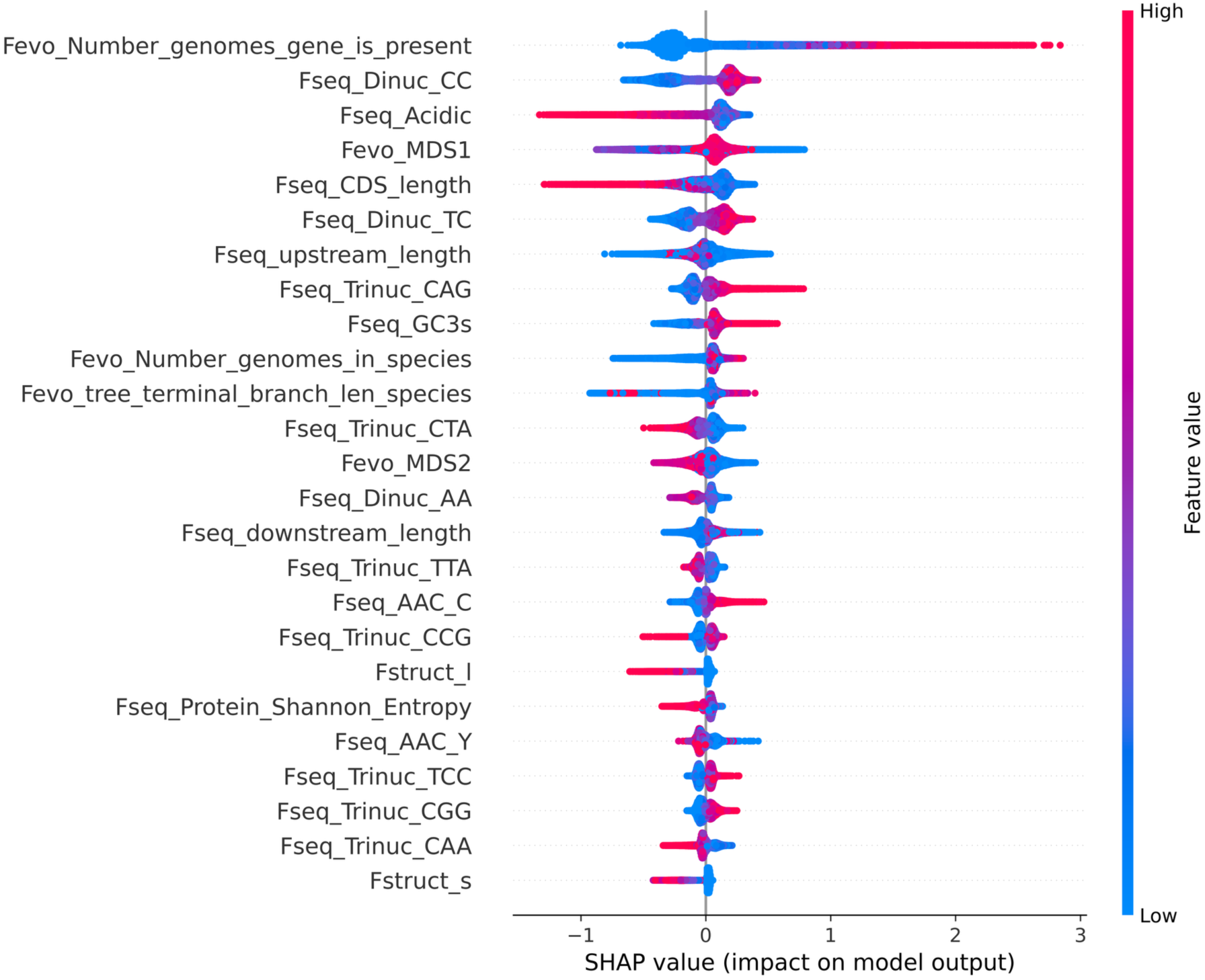
SHAP summary plot showing the top 25 contributing features in the classification of EOGs vs. SOGs with controlling for genomic representation. Positive SHAP values (right side) push predictions toward EOGs; negative values indicate a contribution toward SOGs. Features are colored by their value in the original data (red = high, blue = low). See Table S2 for the complete list.

We found several nucleotide composition and codon usage features, including specific di- and trinucleotide frequencies (e.g., Dinuc CC, Dinuc_TC, Trinuc_CAG) (Figure 5). In addition, SOGs showed more acidic amino acids, suggesting also a distinct amino acid composition compared to EOGs. Additionally, GC content at the third synonymous codon position (GC3s) tends to be higher in EOGs, while coding sequence (CDS) length tends to be higher in SOGs.

**Figure 5.**
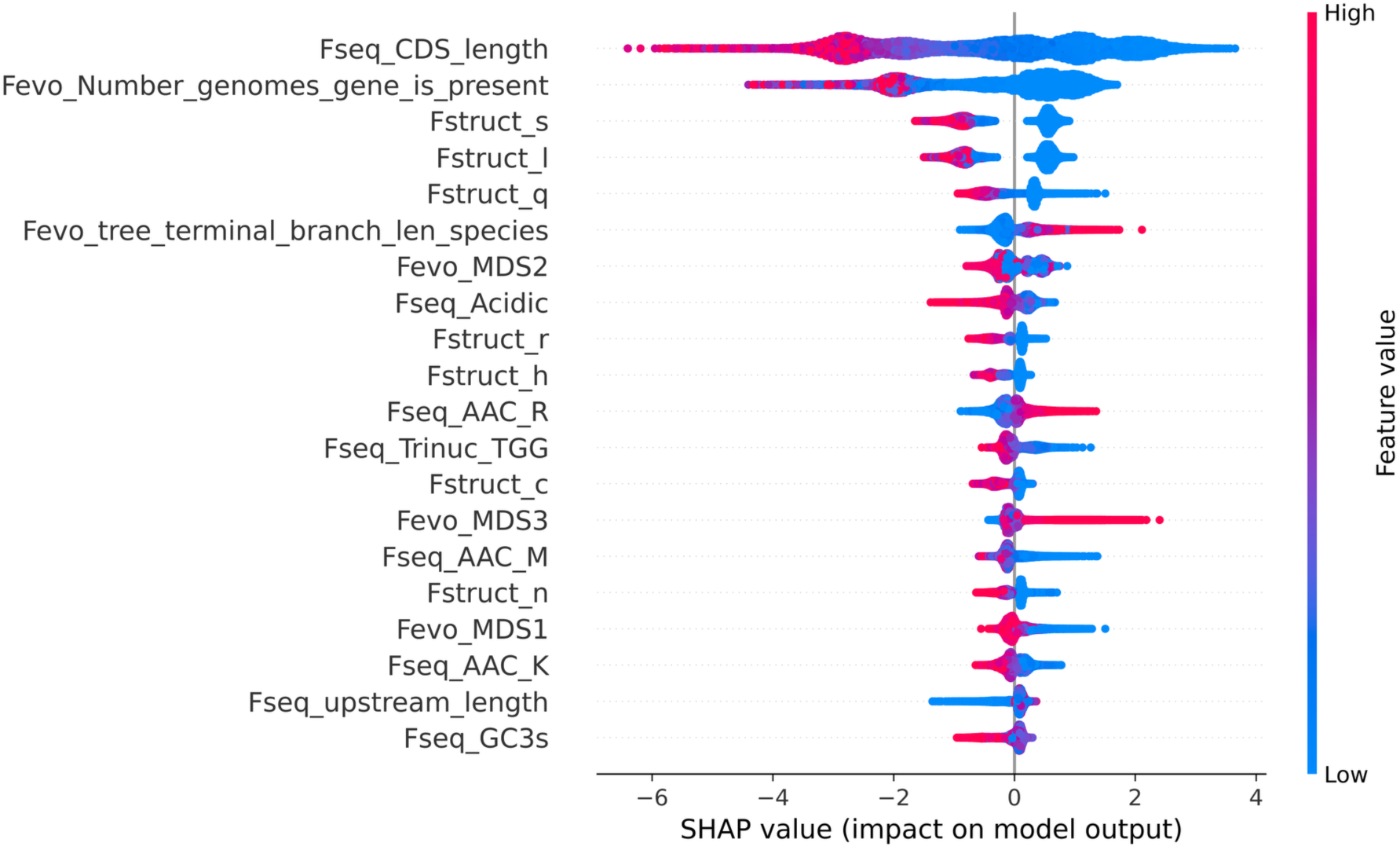
SHAP summary plot showing the top contributing features in the classification of CGs vs. EOGs with controlled genomic representation (Number_genomes_in_species ≥ 700). Positive SHAP values (right side) push predictions toward EOGs; negative values indicate a contribution toward CGs. See Table S3 for the complete list.

We also observed that EOGs are present in a larger number of genomes (higher Number_genomes_gene_is_present values) compared to SOGs, indicating that SOGs represent rarer genes in the pangenomes. We used MDS to reduce the complex pairwise phylogenetic distance relationships from the species tree into a low-dimensional space while preserving the relative evolutionary distances among species. The feature importance of MDS1 and MDS2 suggests that SOGs and EOGs were not uniformly distributed across species, indicating differences in their phylogenetic distribution.

Besides evolutionary features, we also looked at structural features. However, almost no structural features were ranked as important when classifying SOGs and EOGs; only 3Di structural token l and s from ProstT5 appeared among the top 25 features. To validate the SOG and EOG distinction, we considered the analysis of protein-level selection. When genes are spurious, i.e., not coding for real proteins, we would not expect selection on protein-coding genes. Thus, the nonsynonymous (dN) and synonymous substitution (dS) rate would be the same, i.e., dN/dS equals 1. However, dS needs to be larger than 0 to calculate dN/dS and dN/dS estimates were only available for a minority of genes. Among SOGs, only 2,027 out of 26,732 genes (∼8%) had both dN and dS values, whereas among EOGs, 11,500 out of 72,828 genes (∼16%) had such estimates. However, such a small selection of dN/dS values might be biased, which limits the interpretation of the differences in dN/dS between SOGs and EOGs.

### 2.3 Expressed orphan genes differ from conserved genes in sequence, evolutionary, and structural features

Next, we investigated whether the presence of SOGs in the orphan gene category would influence the classification between OGs and CGs. Since we observed a bias by genomic representation, we only include species with at least 700 genomes from now on. We trained machine learning classifiers under two scenarios, using datasets of equal size (18,244 samples per class; 14,595 for training and 3,649 for testing): (1) distinguishing CGs from OGs, and (2) distinguishing CGs from only EOGs. We evaluated model performance using four feature sets after controlling for genomic representation bias (Table 3). We find that model performance improved for sequence, structural, and combined feature sets when classifying CGs vs. EOGs compared to CGs vs. OGs, whereas the evolutionary feature set showed similar performance between the two comparisons.

**Table 3.**
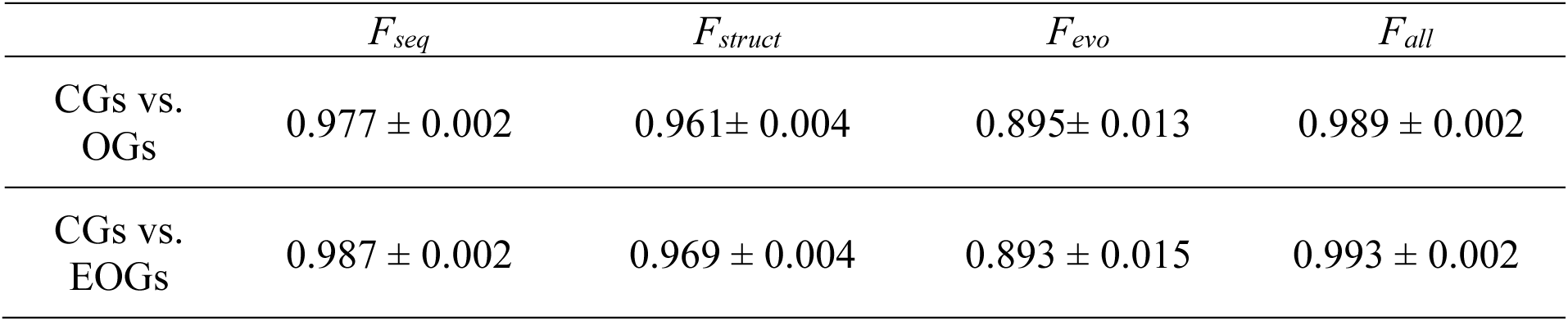
Comparison of classification performance (AUC) for CGs vs. OGs and CGs vs. EOGs using different feature sets, with controlling for genomic representation bias.

We examined the contribution of features to classify EOG and CGs (Figure 5). We focused our interpretation on the top 20 features, as individual feature contributions dropped below 1% after the 20^th^ ranked feature (Figure S5). CDS length emerged as the most important discriminative feature. As shown in the SHAP plot, shorter genes (blue values for CDS length) were strongly associated with EOGs, while longer genes were more likely to be CGs. A similar trend was observed for GC3s, with higher GC3s values occurring more frequently in CGs. Additionally, some evolutionary features are highly important in CGs vs. EOGs comparison. For example, EOGs tend to be present in a lower number of genomes within their species, and this pattern is consistent with the pangenome analysis, where conserved genes were predominantly composed of core genes whereas OGs were largely restricted to the accessory genome. The lower prevalence of EOGs across genomes therefore reflects their tendency to occupy the accessory component of bacterial pangenomes.

Amino acid composition (AAC), including acidic amino acids and the relative abundance of specific residues like arginine (AAC_R), was important to classify EOGs and CGs, suggesting that EOGs encode proteins with distinct biochemical compositions compared to CGs. Moreover, sequence motifs such as trinucleotide frequencies, Trinuc_TGG, appeared among the top features.

We found that several 3Di structural representation features - represented by single-letter tokens (e.g., “a”, “c”, “d”, etc.) extracted via ProstT5 - were highly ranked in feature importance. SHAP analysis of the 3Di features revealed that CGs generally exhibit higher values for several 3Di structural representations, whereas EOGs tend to display lower representation values.

## 3. Discussion

The dataset of microbial orphan genes analyzed here originates from large-scale predictions of species-specific genes in the human gut microbiome (Vakirlis and Kupczok 2024). Various evolutionary processes create orphan genes and bacteria offer large potential to study these due to the ample genomic data that is available at species and strain level (uz-Zaman and Ochman 2025). For example, remodeled gene families have been detected using network approaches (Watson et al. 2022), genes originating from divergence have been distinguished using simulations (Tassios et al. 2026), and de novo genes have been traced using omics approaches (uzZaman and Ochman 2025). Nevertheless, most previous studies have treated all predicted orphans as a single category without considering their evolutionary heterogeneity (Pereira et al. 2025). Additional heterogeneity is created by spurious predictions that can arise from annotation errors or non-functional open reading frames, which has likely obscured the functional and evolutionary signals of true orphans. Here, by explicitly distinguishing between SOGs, which show no expression evidence, and EOGs, we aimed to separate non-expressed artifacts from genuine OGs and to identify the features that best distinguish EOGs vs. SOGs, and EOGs vs. CGs. Our analysis of conserved genes shows that a larger proportion of CGs is expressed compared OGs. This suggests that expression provides a useful signal for distinguishing between *bona fide* and spurious OGs, albeit imperfect for separating more likely functional genes from less likely ones. Our results further show that the apparent importance of evolutionary features is strongly influenced by sampling bias due to the number of genomes representing a species. This pattern is likely explained by the fact that species with more available genomes are typically more abundant in the gut microbiome, and therefore contribute a larger proportion of reads in metatranscriptomic datasets, increasing the likelihood of detecting gene expression (Almeida et al. 2021). Thus, their genomes are better represented in the human gut environment, which could lead to higher expression levels in metatranscriptomics data. To address this, we specifically filtered out species with low genomic representation and restricted the analysis to well-represented species, thereby ensuring that SOGs and EOGs had comparable distributions of genome counts across species. Under this controlled scenario, much of the bias disappeared, highlighting the importance of considering differences between species when studying orphan genes.

Here we analyze feature importance using SHAP values, which jointly evaluate contributions of all features. This approach quantifies not only whether a feature is informative, but also its importance relative to other features, which allows us to analyze all features at once. Feature importance analyses highlight the number of genomes a gene is present in as an informative signal for distinguishing SOGs from EOGs. Many SOGs are restricted to only a few genome occurrences, which are consistent with their likely origin as prediction errors or non-functional sequences. This observation can also be framed in the context of pangenome structure. The pangenome of a species encompasses the complete set of genes across all sequenced genomes and is typically divided into categories. The core genome comprises genes present in nearly all strains, encoding essential or highly conserved functions, whereas the accessory genome contains genes restricted to only a subset of strains (Matthews et al. 2024; Vakirlis and Kupczok 2024). In pangenome studies, such low-frequency genes are known to be enriched in annotation errors or assembly artifacts (Tonkin-Hill et al. 2020). SOGs tend to be rare genes and could thus be also enriched in annotation artefacts. Nevertheless, EOGs were determined using expression signal, which might be easier to detect for high-frequency genes; thus, true low-frequency orphan genes could still be classified as SOGs.

Our approach is able to detect distinguishing sequence features for classifying EOGs vs. SOGs and EOGs vs. CGs. GC content at the third synonymous codon position (GC3s) followed the order CG > EOG > SOG. Given that GC3s is strongly linked to synonymous codon usage bias and is interpreted as a signature of translational selection acting on coding sequences (Wan et al. 2004; Babbitt et al. 2014), the higher GC3s in EOGs relative to SOGs suggests that EOGs are unlikely to be transcriptional artifacts but are instead subject to selective pressures shaping codon usage, potentially enhancing translational efficiency. This is consistent with the even higher GC3s values of CGs indicating stronger selection in conserved genes. From an evolutionary perspective, the intermediate GC3s levels observed in EOGs suggest that OGs are relatively young coding sequences that are in the process of adapting to the cellular environment and gradually acquiring functional characteristics. In contrast, the lower GC3s values in SOGs are consistent with weak or absent selective constraints on codon usage, as expected for non-functional open reading frames. Beyond GC3s, we observed that several dinucleotides show differential enrichment between EOGs and SOGs, but not between EOGs and CGs. Dinucleotide composition is shaped by genome-wide mutational processes and can serve as a genomic signature that characterize the sequence composition of genes (Kariin and Burge 1995; Chen et al. 2014). Thus, SOGs show sequence compositions that deviate from EOGs consistent with the expectation that SOGs represent spurious ORFs or annotation artifacts. Furthermore, CDS length emerged as one of the most informative features across comparisons. SHAP analysis revealed that: CGs possess longer CDSs than EOGs, while SOGs also tend to have longer CDSs than EOGs. The short length of EOGs is consistent with their potential role as recently emerged genes (Tautz and Domazet-Lošo 2011; Vakirlis and Kupczok 2024). The longer CDSs observed in SOGs, despite lacking expression support, likely result from annotation artifacts. Gene prediction pipelines are biased toward reporting long open reading frames (Choudhary et al. 2020), for example, gene-prediction pipelines such as Prokka (Seemann 2014) and Prodigal (Hyatt et al. 2010) tend to detect long open reading frames, thereby increasing the chance of spurious annotations in sequences without experimental support. Thus, SOGs may appear long in sequence, but this does not equate to functional or structural validity.

*De novo* genes in eukaryotes are often thought to originate from previously non-coding open reading frames (ORFs), but in prokaryotic genomes, which are highly compact and dense in coding regions, new genes may more commonly arise from alternative reading frames within existing coding sequences (Watson et al. 2022). Thus, some de novo–originated EOGs likely arose from alternative reading frames. Our results have shown that EOGs exhibit different proportions of amino acids compared with CGs and SOGs. Amino acid composition is a major determinant of intrinsic structural disorder (Wilson et al. 2017), and is therefore subject to biophysical and evolutionary constraints. A previous study reported that predicted intrinsic structural disorder is highest in de novo proteins, intermediate in random sequences, and lowest in conserved proteins (Middendorf and Eicholt 2024). Although structural disorder does not appear as an important feature in any of our analyses, the observed differences in amino acid composition, particularly in the content of acidic amino acids, across EOGs, SOGs, and CGs could reflect differences in the strength of selection acting on these genes. CGs encode proteins shaped by long-term selection, whereas EOGs likely represent younger proteins that have begun to acquire functional constraints. In contrast, SOGs, which lack expression support, are expected to be under little or no selective pressure and may therefore exhibit amino acid compositions that are closer to those of random sequences.

Interestingly, only few structural features appeared among the top predictors in our SHAP analysis for distinguishing EOGs from SOGs. This likely reflects a methodological limitation rather than the absence of structural signals. Specifically, the structural predictor we used was ProstT5 (Heinzinger et al. 2024), which itself was trained on AlphaFold’s protein sequences and 3Di information, and the 3Di representation was obtained via Foldseek (van Kempen et al. 2024). Because AlphaFold relies on multiple sequence alignments, its structural embeddings are not independent of evolutionary conservation information. Consequently, structural features may overlap with the sequence features already used in the model, reducing their apparent contribution to orphan genes prediction.

In summary, our analysis shows that a substantial proportion of predicted orphan genes in gut microbiomes are likely spurious, while a smaller subset exhibits consistent expression. By integrating expression evidence with sequence, structural, and evolutionary features, we established a framework for the discrimination between true orphan genes and prediction artifacts. The findings enhance our understanding of orphan genes and provide a foundation for future studies on the origination and functional roles of orphan genes in microbial communities.

## 4. Materials and Methods

### 4.1 Dataset construction

In a previous study, orphan genes (OGs) were identified through a multi-step filtering process (Vakirlis and Kupczok 2024). First, the Unified Human Gastrointestinal Protein catalogue, clustered into protein families at 50% identity (UHGP50 v1.0) (Almeida et al. 2021) was analyzed and protein families present in multiple species were removed, along with those having a match in EggNOG (Buchfink et al. 2021; Hernández-Plaza et al. 2023). Next, DIAMOND searches were performed against the NCBI prokaryotic RefSeq database, removing sequences with significant matches (E-value < 10^-3^), except for those with >90% coverage and >95% identity, which were likely from the same species. A final BLASTp (Altschul 1997) search against the UHGP50 protein catalogue further filtered out sequences matching proteins from different species (E-value < 10^-5^). To reduce contamination, OGs found on contigs containing only orphans were removed. The remaining sequences formed our reference dataset of 631,104 species-specific OGs, while the non-orphan families were labelled conserved genes (CGs), totaling 4,104,442. In the following, we focused on highly conserved genes (defined as genes present in >10 species, 26,545 genes in total) rather than all conserved genes because their broad phylogenetic distribution provides stronger evidence of long-term evolutionary conservation, making them a more reliable group to compare to orphan genes.

The Snakemake (v7.32.4) (Mölder et al. 2021) workflow management system was used to develop an automated pipeline (Figure 1, https://github.com/cheche0109/OGs-SOGs-CGs). The pipeline includes processing and alignment of metatranscriptomics sequencing data sets against our orphan reference dataset and generating BAM files for further analysis to distinguish EOGs and SOGs.

#### 4.1.1 Search and download of metatranscriptomics data

We analyzed metatranscriptomics data to label expressed genes in our reference dataset as EOGs, and non-expressed genes as SOGs. To this end, we searched for libraries containing raw metatranscriptomics data in the sequence read archive (SRA) (Katz et al. 2022) (10 April 2024) database of the National Center for Biotechnology Information (NCBI). To ensure that the included metatranscriptomics data originated from prokaryotic human gut metagenomes, we used the following criteria: (i) human gut metagenome, NOT mouse gut metagenome, (ii) AND metatranscriptomics, AND biomol RNA, (iii) AND prokaryote, NOT virome. The compressed RNA-seq read data (.fastq.gz) of the accessions was downloaded directly using fasterq-dump (v3.1.1) (Leinonen et al. 2011) from the SRA Toolkit.

#### 4.1.2 Read processing

Cutadapt (v4.0) (Martin 2011) with the parameter -q 30 was used to trim low-quality bases from the compressed sequencing reads downloaded from the SRA database. Trimmed reads were mapped to our orphan reference genes with CoverM (v0.7.0) (Aroney et al. 2025) using the command *coverm make --discard-unmapped -p bwa-mem2*, generating BAM files directly converted from the SAM outputs. Reads with a mapping quality score of 0 were then removed using the command *samtools view -hbq 1* to eliminate low-confidence alignments and ensure accurate coverage estimation.

#### 4.1.3 Labeling EOGs and SOGs

We used CoverM with the command *coverm contig -- min-read-percent-identity 95 --min-read-aligned-percent 75 --min-covered-fraction 0.6 -- contig-end-exclusion 0* to calculate read coverage per gene from the bam files, without excluding any base pairs from the beginning or end of each gene. To ensure that only highly similar reads are mapped, we require at least 95% identity to the reference gene and to exclude short, ambiguous matches, we require a minimum of 75% of the read length aligned to the gene. Finally, to ensure that the detected expression is biologically meaningful rather than resulting from fragmented or low-confidence mappings, we retained only genes with at least 60% of their length covered by mapped reads. Genes that did not meet the defined thresholds for any library were classified as SOGs. For the remaining genes, we explored the influence of threshold settings on coverage and the number of libraries in which a gene was detected. Genes exceeding the coverage threshold and present in at least 2 libraries were designated as EOGs.

#### 4.1.4 Expression analysis of CGs

To evaluate whether the absence of detectable expression is a reasonable criterion for identifying potentially SOGs, we performed a parallel expression analysis on conserved genes (CGs). To reduce bias caused by uneven genomic representation across species, we restricted the analysis to species with Number_of_genomes_in_species ≥ 700, consistent with the filtering strategy used in our orphan gene analyses. We then estimated low, medium, and high genomic representation levels using the 25%, 50%, and 75% quantiles of Number_of_genomes_in_species among the eligible species. Based on these thresholds and the species phylogeny, we selected three phylogenetically distant representative species with different genome representation levels: GUT_GENOME001602 (*Holdemanella* sp002299315), GUT_GENOME239670 (*Bifidobacterium pseudocatenulatum*), and GUT_GENOME140299 (*Prevotella copri*).

For each selected species, we extracted CGs belonging to that species by retaining genes present in more than 10 species across the UHGP-50 dataset. Expression analysis was performed using the same Snakemake pipeline and the same set of 4,969 human gut metatranscriptomic libraries described above. Read processing, mapping, and coverage estimation were conducted identically to the procedure used for OGs. A CG was considered expressed if it showed detectable coverage in the metatranscriptomic data according to the same criteria applied to OGs.

### 4.2 Feature construction

Several features were derived from our previous study (Vakirlis and Kupczok 2024), including:

i. **Sequence features** — GC content, GC content at the third synonymous codon position, biosynthetic cost, isoelectric point, coding sequence (CDS) length, CpG dinucleotide frequency, and protein self-aggregation propensity (length and energy);
ii. **Structural features** — number of predicted transmembrane domains, percentage of transmembrane regions, and percentage of intrinsically disordered regions;
iii. **Evolutionary features** — number of genomes in which the gene is detected, total number of genomes available for the corresponding species, and terminal branch length for each species, taken from the phylogenetic tree of Almeida et al. (Almeida et al. 2021).

We included additional features for the three groups as follows.

#### 4.2.1 Sequence features

We processed DNA and protein fasta files to extract sequence-based features. For DNA sequences, we computed Shannon entropy, dinucleotide frequencies, and trinucleotide frequencies. For protein sequences, we calculated amino acid composition (AAC) and physicochemical properties with Biopython (version 1.83), including Kyte-Doolittle hydropathy scores (to evaluate protein hydrophobicity), average residue weight, and average charge at pH 7.0. Additionally, amino acids were classified (tiny, small, aliphatic, aromatic, non-polar, polar, charged, basic, and acidic) based on their size, structure, and charge, using Pepstats from the EMBOSS tools (Rice et al. 2000). The presence of signal peptides was predicted using SignalP 6.0 (Teufel et al. 2022) in fast mode. Each sequence was processed individually, and the extracted features were stored in a CSV file for downstream analysis.

#### 4.2.2 Structural features

Protein secondary structure was predicted using the RaptorX Predict_Property package (v1.01) in default mode, and the percentage composition of disordered regions, helices, and strands was computed. Additionally, the protein language model ProstT5 (Heinzinger et al. 2024) was used to translate 1D amino acid sequences into 3Di structural representations. Based on the T5 transformer architecture, ProstT5 was implemented using Hugging Face’s Transformers library and executed on CUDA-enabled hardware. The model output was stored in FASTA format, and the percentage composition of the 20 distinct 3Di structural representations per sequence was calculated.

#### 4.2.3 Evolutionary features

We first computed the pairwise phylogenetic distance matrix for all bacterial species based on their positions in the reference phylogenetic tree provided in newick format, as computed by Almeida. The distance between each species pair corresponds to the total branch length separating them on the tree, as calculated using the ETE3 toolkit (Huerta-Cepas et al. 2016), reflecting the evolutionary divergence among species. The computed pairwise distances were then projected into a three-dimensional space using metric multidimensional scaling (MDS) implemented in scikit-learn (Buitinck et al. 2013) (version 1.3.2).

To assess the discriminative power of different feature sets in classifying CGs, SOGs, EOGs, we evaluated the effectiveness of various feature groups. Specifically, we trained classification models using 4 feature sets:

1. *F_seq_* – 122 sequence features.
2. *F_struct_* – 26 structural features.
3. *F_evo_* – 6 evolutionary features.
4. *F_all_* – 154 features, combining *F_seq_*, *F_struct_*, and *F_evo_*.

### 4.3 Machine learning

To classify SOGs versus EOGs, and EOGs versus CGs, we developed machine learning models based on XGBoost (Chen and Guestrin 2016). For each comparison, we constructed a balanced dataset by randomly downsampling the majority class to achieve equal representation of both classes. Additionally, prior to model training, z-score normalization was applied using parameters (mean and standard deviation) estimated from the training set only, which were then used to normalize the test data.

Classifiers were trained on *F_seq_*, *F_struct_*, *F_evo_*, and *F_all_*. Model selection and hyperparameter tuning were performed using a nested cross-validation strategy: an inner 4-fold cross-validation for hyperparameter optimization, and an outer 5-fold stratified cross-validation to assess generalization performance. The hyperparameter grid search explored the following parameter ranges: n_estimators: [100, 200], max_depth: [3, 5, 7], learning_rate: [0.005, 0.01, 0.1], subsample: [0.8, 1.0], colsample_bytree: [0.8, 1.0].

Model performance was primarily evaluated using the area under the receiver operating characteristic curve (AUC). For each comparison, we report the mean AUC ± standard deviation (SD) across inner folds. SD reflects variability among the folds. For interpretability, we extracted feature importance from the trained models and applied SHAP (SHapley Additive exPlanations) (Lundberg and Lee 2017) to quantify the contribution of each feature to the classification outcome.

## Data Availability

The data underlying this study are provided within the manuscript and its Supplementary Information files. The complete Snakemake pipeline used for data processing and model training is available at https://github.com/cheche0109/OGs-SOGs-CGs. Metatranscriptomic data were retrieved from the NCBI Sequence Read Archive (SRA) (https://www.ncbi.nlm.nih.gov/sra), and the full list of SRA accession numbers used in this study is available at https://github.com/cheche0109/OGs-SOGs-Gs/blob/main/metatranscriptomics/runs_4969.txt.

