## Supplementary material for "Identification and Classification of Expressed Orphan Genes, Spurious Orphan Genes, and Conserved Genes in the Human Gut Microbiome": Figure S

##### Feature groups

###### *$F_{seq}$: 122 Sequence-Based Features*

- **Nucleotide composition:** GC content, GC3s, CpG dinucleotide frequency, and coding sequence length (CDS_length).
- **Dinucleotide and trinucleotide motifs:** Frequencies of all 16 possible dinucleotides (e.g., Dinuc_AT, Dinuc_CG) and all 64 possible trinucleotides (e.g., Trinuc_AAA, Trinuc_GTG).
- **Sequence complexity and property:** DNA and protein Shannon entropy, isoelectric point, biosynthetic cost, average hydropathy, average residue weight, and average charge.
- **Amino acid property:** Proportions of tiny, small, aliphatic, aromatic, non-polar, polar, charged, basic, and acidic residues, as well as frequency of individual amino acids (AAC_A, AAC_C, AAC_D, etc.). Presence of signal peptides (signalp), PASTA aggregation length and energy.
- Genome property: upstream/downstream region lengths,

###### *$F_{struct}$: 26 Structural Features:*

- **Secondary structure content:** Percentage of residues in coil (C_Percentage), helix (H_Percentage), and strand (E_Percentage) conformations.
- **Transmembrane and disorder predictions:** Number of predicted transmembrane domains (PHOBIUS_no_tm_domains), predicted proportion of disordered regions (PredProp_diso_pct), and transmembrane regions (PredProp_tm2_pct).
- **3Di representation from ProstT5:** a,c,d,e,f,g,h,i,k,l,m,n,p,q,r,s,t,v,w,y

###### *$F_{evo}$ : 6 Evolutionary Features*

- **Gene presence:** Number of genomes in which the gene is present (Number_genomes_gene_is_present), and the total number of genomes per species (Number_genomes_in_species).
- **Phylogenetic branch length:** Terminal branch length of the gene tree within a species (tree_terminal_branch_len_species).
- **Multidimensional scaling (MDS):** Three coordinates (MDS1, MDS2, MDS3) reflecting evolutionary or gene similarity distances.

#### Supplementary figures

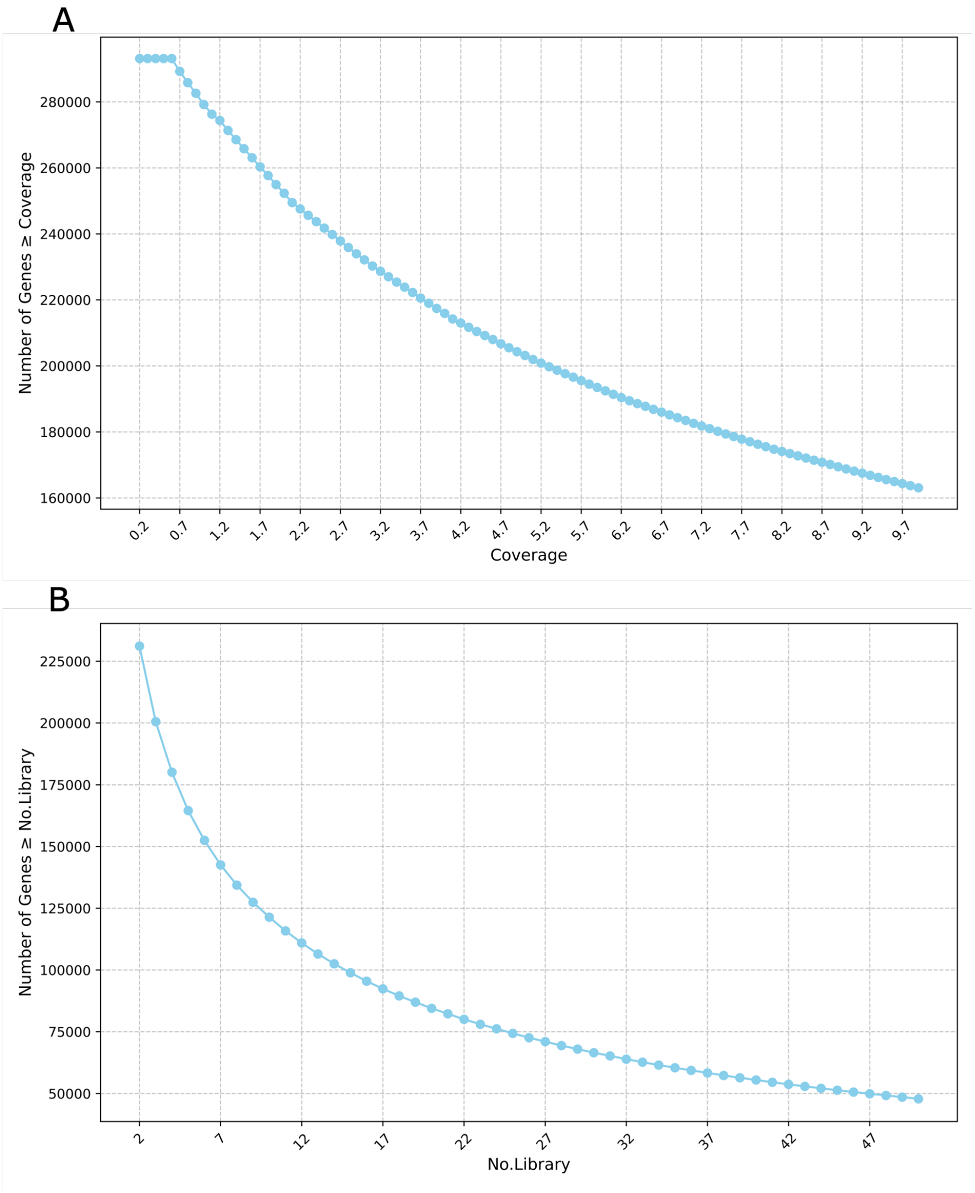


**Figure S1** (A) The number of reference genes detected in at least one of the 4,969 prokaryotic human gut metatranscriptomics libraries, shown across varying coverage thresholds. The x-axis represents the coverage threshold, the y-axis the number of reference genes meeting or exceeding the given threshold. (B) The number of reference genes detected across varying numbers of prokaryotic human gut metatranscriptomics libraries. The x-axis represents the number of libraries included, and the y-axis shows the number of reference genes that are detected in at least that many libraries (i.e., genes meeting or exceeding the given library-presence threshold).

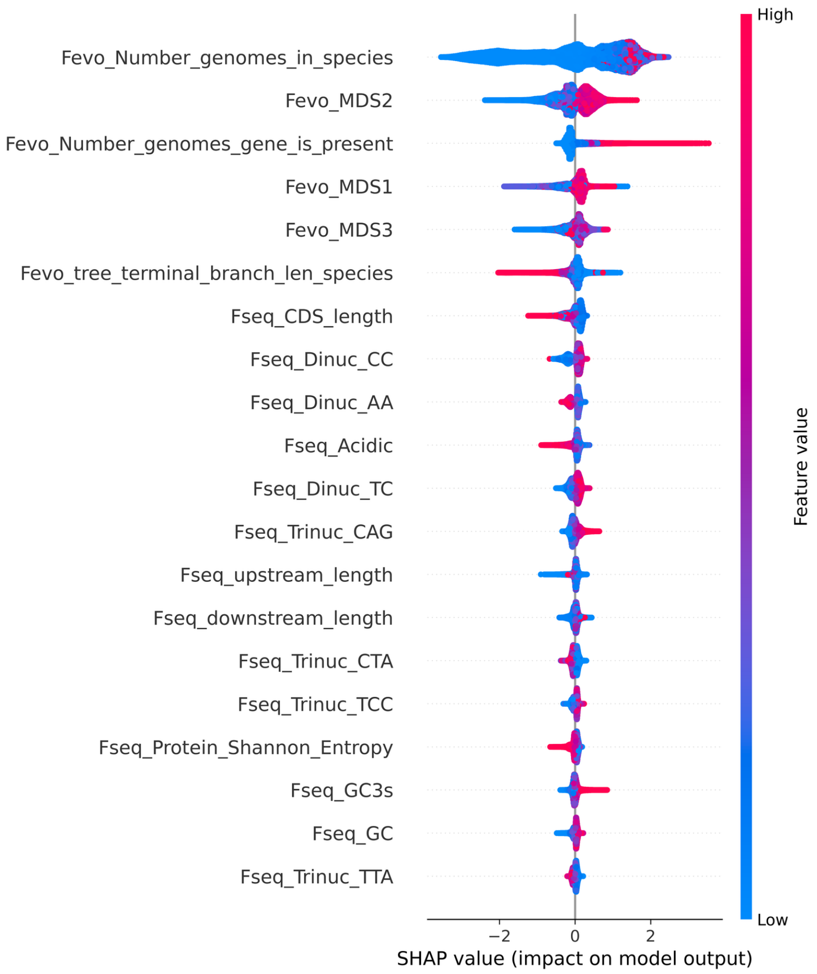


**Figure S2.** SHAP summary plot showing the top 20 contributing features in the classification of EOGs vs. SOGs without controlling genomic representation. Positive SHAP values (right side) push predictions toward EOGs; negative values indicate a contribution toward SOGs.

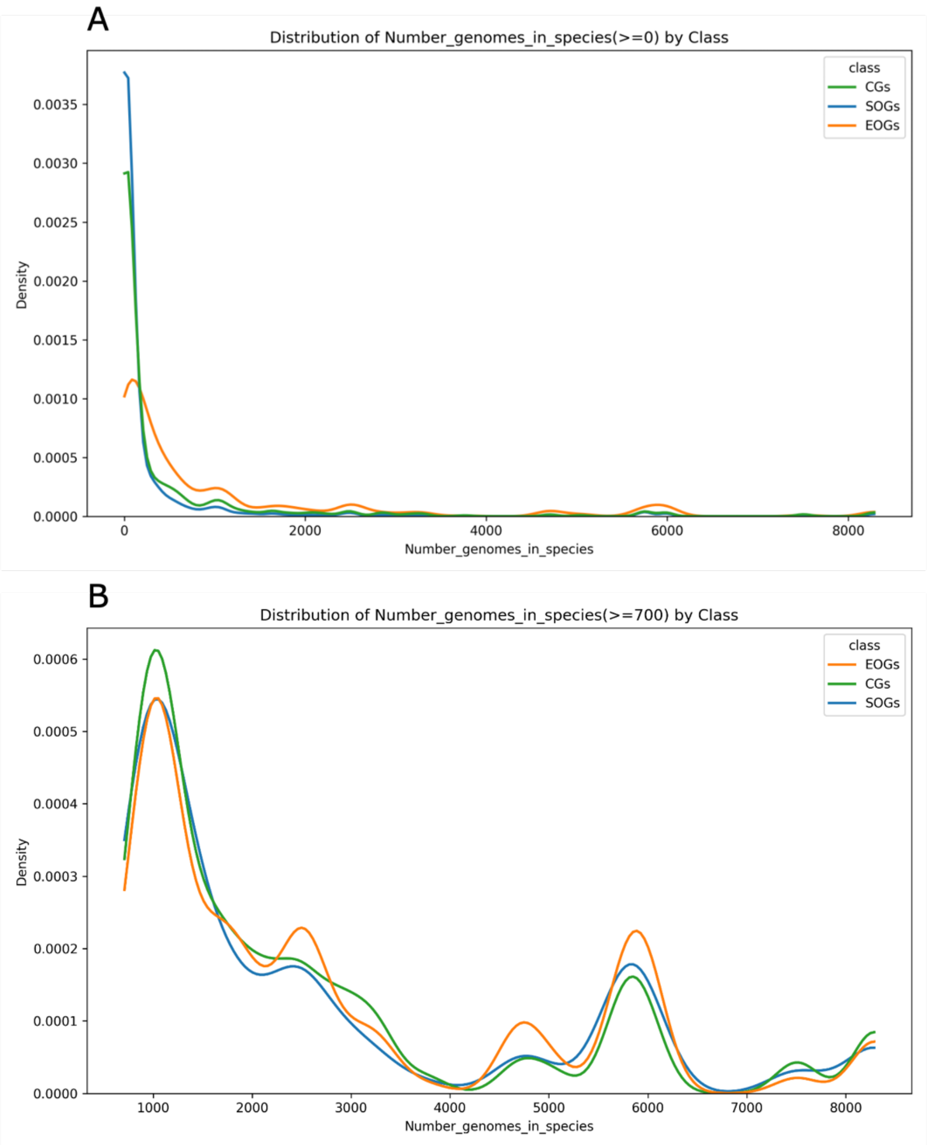


**Figure S3.** Distribution of the total number of genomes available in species per gene. (A) Distribution across all species (no filtering). (B) Distribution after filtering for genomic representation, i.e., species with at least 700 genomes (Number_genomes_in_species ≥ 700).

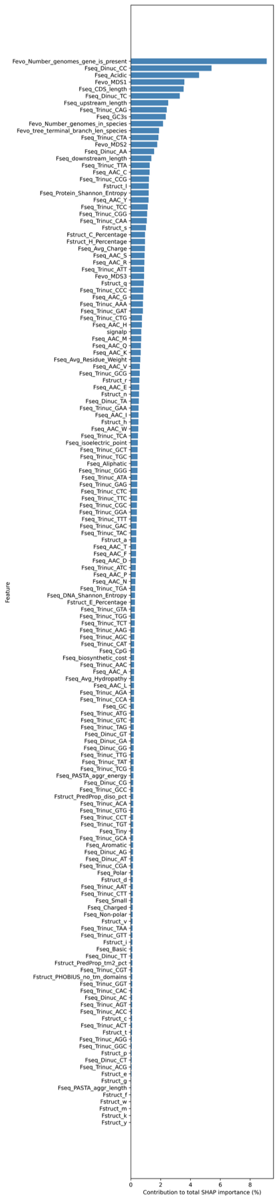


**Figure S4**. Feature importance for EOGs versus SOGs and individual feature contribution based on average absolute SHAP values.

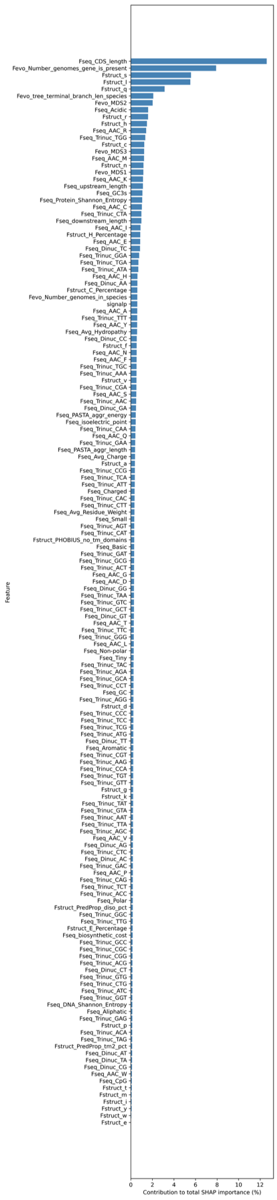


**Figure S5.** Feature importance for EOGs versus CGs and individual feature contribution based on average absolute SHAP values.

### Supplementary tables

Table S1: List of 88 species after controlling the genomic representation bias

Table S2: SHAP values for all features from classification EOGs vs. SOGs (after correcting for genomic representation)

Table S3: SHAP values for all features from classification EOGs vs. CGs (after correcting for genomic representation)
